# Identification and characterization of a small-molecule inhibitor of the *Pseudomonas aeruginosa* SOS response

**DOI:** 10.1101/2025.05.13.651886

**Authors:** Filippo Vascon, Benedetta Fongaro, Vytautas Mickevičius, Antonella Pasquato, Birute Grybaite, Vidmantas Petraitis, Patrizia Polverino de Laureto, Donatella Tondi, Povilas Kavaliauskas, Laura Cendron

## Abstract

The SOS response is among the most conserved pathways that promote the acquisition of antibiotic resistance in bacteria. This study aimed to identify and characterize small-molecule inhibitors of the SOS response system in the opportunistic pathogen *Pseudomonas aeruginosa*. A library of 318 drug-like compounds was screened for inhibition of RecA-induced LexA autoproteolysis, a key step in SOS activation. One hit compound, 3-(2-sulfanylanilino)propanoic acid, showed dose-dependent inhibition with an IC_50_ in the mid-micromolar range. Differential scanning fluorimetry and isothermal titration calorimetry analysis revealed that A12 binds to both RecA and LexA with low micromolar affinity. Mass spectrometry analysis demonstrated that A12 covalently modifies RecA likely via condensation, while it forms a disulfide bond with Cys104 of LexA. Inhibition was diminished under reducing conditions, confirming disulfide formation is crucial for A12 activity. Importantly, A12 did not impair LexA’s ability to bind SOS box DNA sequences, which is needed to keep the SOS genes repressed. While A12’s potency requires optimization, it represents a promising scaffold for developing anti-SOS compounds targeting *P. aeruginosa*.

## Introduction

*Pseudomonas aeruginosa* is an opportunistic bacterial pathogen of high clinical relevance, characterized by a broad spectrum of resistance to currently available antibiotics [1]. Finding new therapeutic strategies that efficiently and selectively kill this microorganism or increase its sensitivity to antibiotics is thus a priority.

The bacterial SOS response to DNA damage regulates DNA repair and mutagenesis, thus representing a driver of antibiotic resistance. Moreover, in *P. aeruginosa* this pathway controls the expression of virulence factors and the formation of biofilms, so its inhibition could be greatly beneficial for the eradication of *P. aeruginosa* infections.

We and others have recently solved the long-standing question on how the two SOS response master regulators (namely the DNA-damage sensor RecA and the autoproteolytic transcriptional repressor LexA) interact with each other, showing a high structural conservation between *E. coli* and *P. aeruginosa* [2–4].

The druggability of the SOS response by small molecules has already been demonstrated, but no anti-SOS compounds has entered clinical testing. The SOS system presents several hurdles that have likely hindered the discovery of a potent inhibitor with drug-like properties, including: (*i*) the homology of RecA with the human Rad51 family, that calls for cautious off-target evaluation, (*ii*) the limited dimensions and strong substrate specificity of LexA active site, which reduce the explorable chemical space of potential inhibitors, and (*iii*) the intramolecular nature of LexA proteolysis (the core of the SOS activation), which implies an effective local substrate concentration hard to be overwhelmed by ectopic competitive inhibitors.

These challenges were recently overcome by anti-LexA nanobodies that target LexA cleavage site and its conformational dynamics [5], but this approach still needs optimization, in particular in terms of delivery routes.

In this context, new high-throughput screenings of molecular fragments could provide promising scaffolds for suppressing the SOS response via previously unexplored pharmacophores and mechanisms of action.

The highest advantages of small, drug-like compounds over therapeutic peptides and biomolecules are mainly related to their membrane permeability, pharmacokinetic profile and potential stability. In particular, their low molecular weight (< 1 kDa) and usually high hydrophobicity allow them to easily diffuse through biological membranes, reaching cytoplasmic targets (as are RecA and LexA in bacteria).

The objective of studies reported in this work is to identify and characterize potential inhibitors of *P. aeruginosa* SOS response among a library of drug-like compounds, which has been previously screened and provided inhibitors of proteolytic enzymes, such as Furin [6].

A hit compound (hereafter referred to as *A12*) emerged from the primary library screening and its effect on *P. aeruginosa* SOS system (RecA_Pa_ and LexA_Pa_) was extensively analyzed *in vitro* by LexA autoproteolysis inhibition assays, differential scanning fluorimetry, isothermal titration calorimetry, electrophoretic mobility shift assay and mass spectrometry. A12 confirmed the ability to inhibit LexA_Pa_ self-cleavage promoted by activated RecA_Pa_ (RecA_Pa_/ssDNA/ATP S, *RecA*_*Pa*_*\**), with an IC_50_ in the mid-micromolar range. Mechanistic dissection evidenced a dual targeting which requires further elucidation and the examination of A12-based libraries of compounds that explore a broader chemical space.

## Results

### Screening of a library of small molecules and validation of selected hits

A custom made library of 318 small molecules (MW < 1000 Da) has been screened at two different concentrations (25 µg/mL and 125 µg/mL, corresponding to molar concentrations of 30-170 µM and 150-850 µM, respectively) on the *P. aeruginosa* SOS system by a previously validated fluorescence polarization (FP)-based RecA*-induced LexA autoproteolysis *in vitro* assay [4,5,7– 9]. 1 µM FlAsH-LexA_Pa_^CTD^ and 1 µM pre-activated RecA*_Pa_ were incubated with the screened chemical fragments 1h at 37 °C before measuring FP.

Two compounds (referred to as “A12” and “E17”) displayed an increase of LexA autoproteolysis inhibition reasonably consistent with the tested concentrations and exceeded 50% inhibition at 125 µg/mL (Fig. 1 A).

**Fig. 1:**
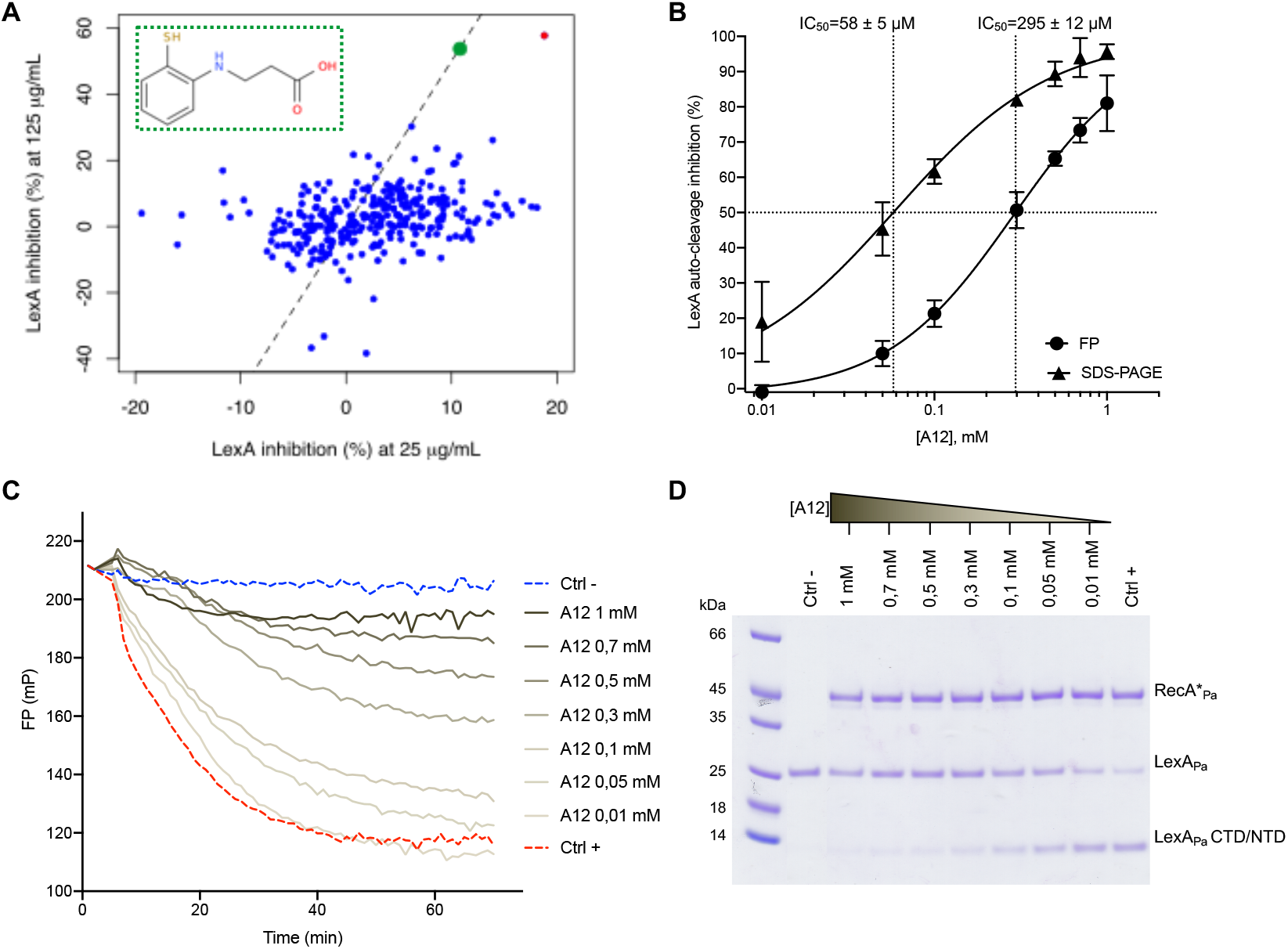
In-vitro identification of a new inhibitor of *P. aeruginosa* SOS system. (A) LexA autoproteolysis percent inhibition values obtained in the initial FP-based library screening at two concentrations (25 and 125 µg/mL). Among the two compounds showing a consistent activity at the two concentrations tested, one revealed to be a false positive (red dot) while compound A12 was a true positive (green dot and inset). The dotted line represents y=5x function. (B) Dose-response curves of A12 on RecA*_Pa_-induced LexA_Pa_ autoproteolysis, as obtained by FP (one replicate shown in panel C) and SDS-PAGE (one representative gel shown in panel D) assays. Data points in panel B represent averages of 3 replicates ± standard deviation. Uncertainties on IC_50_ values correspond to standard error.

Then, the FP-based LexA autoproteolysis assay was repeated on the two hits, monitoring the full kinetics of LexA cleavage over a 1-hour incubation and exploring a wide range of drug concentrations (10-1000 µM). Compound E17 proved to be a false positive as it induced evident protein aggregation events (Fig. S1), causing a very high FP increase after adding the compound to the protein mixture.

Conversely, A12 manifested a negligible protein precipitation effect and a clear dependence of LexA inhibition on drug concentration, reaching 80% of LexA self-cleavage suppression at 1 mM. Higher concentrations of A12 were not explored since its solubility in aqueous buffers strongly worsened above 500 µM and would have misled any inhibition data.

An IC_50_ of 295 ± 12 µM was obtained from data fitting (Fig. 1 B-C). A12 was disclosed to be 3-(2-sulfanylanilino)propanoic acid (structure in Fig. 1 A, inset).

To further validate the activity of A12 on RecA*_Pa_-LexA_Pa_, an assay was performed using full-length LexA_Pa_ and autoproteolysis was followed by SDS-PAGE in the presence of different concentrations of A12 (Fig. 1 D). Cleavage reactions were performed at 1 µM of full-length LexA_Pa_, initiated by the addition of 1 µM RecA*_Pa_, and stopped after a 1-hour incubation at 37 °C, corresponding to the same conditions used for the FP assay.

SDS-PAGE analysis of LexA_Pa_ autoproteolysis demonstrated a clear reduction of cleavage products (and a corresponding intensification of the band of uncleaved LexA_Pa_) at increasing concentrations of A12. Data fitting resulted in an IC_50_ of 58 ± 5 µM, which is 5 times lower than the value obtained in the FP assay. This discrepancy might result from intrinsic differences and limitations of the two techniques, or from a higher affinity of A12 to the full-length LexA_Pa_ than its CTD domain alone.

### A12 binding to SOS response players

To elucidate whether A12 binds RecA_Pa_ or LexA_Pa_, the two proteins were submitted to a thermal shift assay by differential scanning fluorimetry (DSF), in the presence of different concentrations of A12 (30-1000 µM; Fig. 2 A-C). Controls devoid of A12 were included as well. To avoid any potential self-cleavage of LexA_Pa_, the S125A inactive mutant was used.

**Fig. 2:**
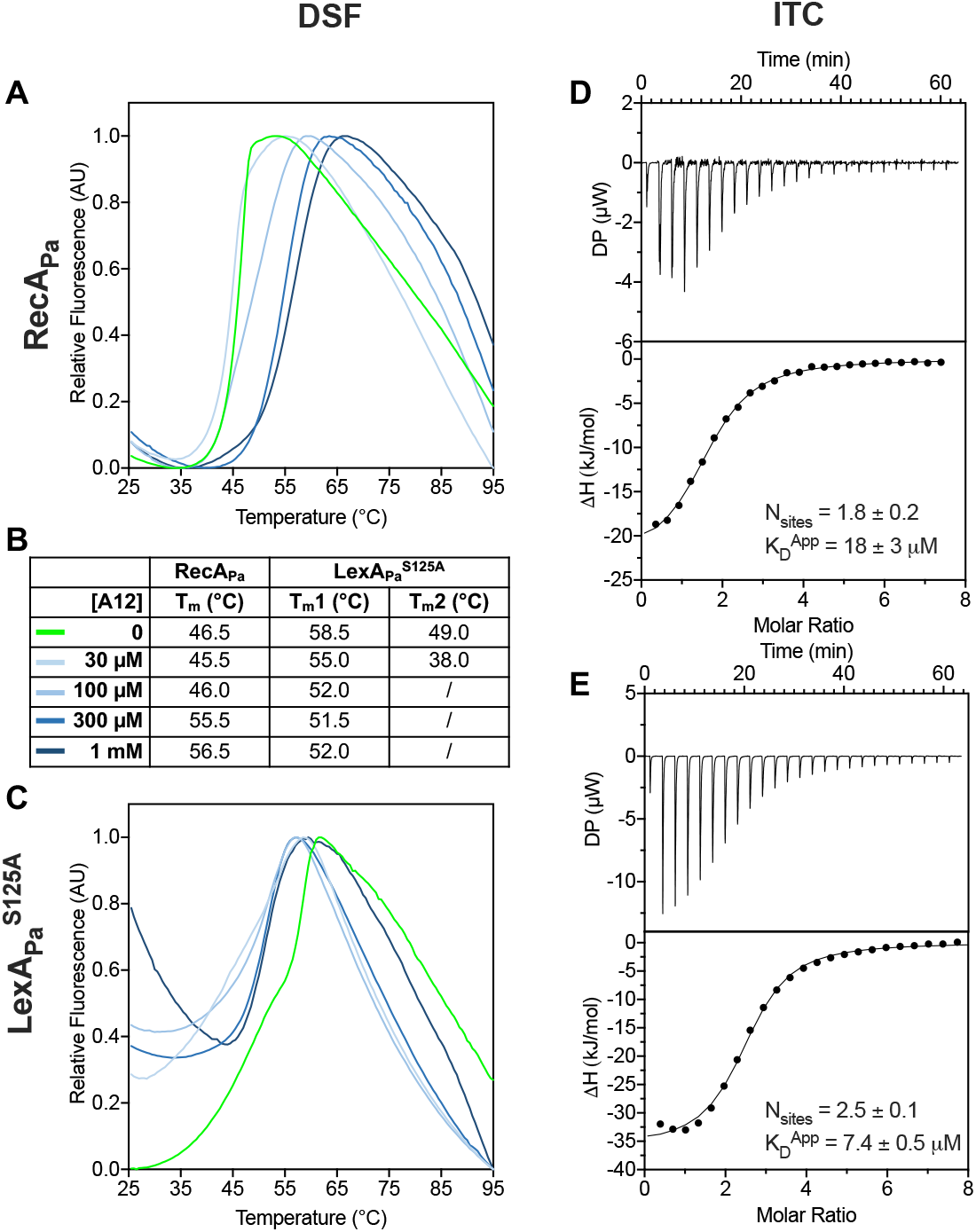
Analysis of A12 binding to RecA_Pa_ and LexA_Pa_^S125A^ by Differential Scanning Fluorimetry (DSF) and Isothermal Titration Calorimetry (ITC). (A, C) Normalized melting curves of RecA_Pa_ and LexA_Pa_^S125A^ in the presence of different concentrations of A12. One representative of two coherent replicates is reported. (B) Melting temperatures, corresponding to local or global minima of melting curve derivatives. (D,E) Raw traces (top) and integrated binding isotherms (bottom) of ITC analysis of A12 on purified RecAPa and LexA_Pa_^S125A^ at 298 K. Integrated heats were fitted by a «one set of sites» model. One representative curve is shown for each condition. The number of equivalent binding sites (N_sites_) and the apparent equilibrium dissociation constant (K_D_) are represented as average ± S.D., calculated on at least two replicates.

The sharp melting curve of RecA_Pa_ (T_m_ 46.5 °C) was noticeably shifted rightwards by increasing concentrations of A12, with a significant ΔT_m_ of 10 °C at 1000 µM A12 compared to the control (Fig. 2 A-B), a clear indication of A12 binding to RecA_Pa_.

Concerning full-length LexA_Pa_^S125A^, a biphasic melting curve was obtained for the pure protein (T_m1_ 49.0 °C; T_m2_ 58.5 °C; Fig. 2 C), most likely due to the independent denaturation of its two domains (the DNA-binding NTD and the autoproteolytic CTD) assembled into dimers. Treatment with A12 at increasing concentrations induces some unexpected events on LexA_Pa_^S125A^: at 30 µM, A12 seems to cause a general destabilization of LexA_Pa_^S125A^, as both the melting transitions are shifted towards lower temperatures, while in the presence of A12 above 100 µM monophasic melting curves are obtained, with a flexus between T_m1_ and T_m2_ (at 52.0 °C; Fig. 2 B). This behavior suggests that A12 is inducing some conformational rearrangement on LexA_Pa_^S125A^, probably interacting with a region at the boundary between the two domains or close to the dimerization interface.

To investigate deeper the binding of A12 to the two members of *P. aeruginosa* SOS complex, Isothermal Titration Calorimetry (ITC) was performed on RecA_Pa_ and LexA_Pa_^S125A^ (Fig. 2 D-E). The integrated binding isotherms of both proteins were best fitted by a “one set of sites” model. Roughly two equivalent binding sites were detected on each protein, while the affinity of A12 for LexA_Pa_^S125A^ (*K*_D_ = 7.4 ± 0.5 μM) was slightly higher than for RecA_Pa_ (*K*_D_ = 18 ± 3 μM). Taken together, DSF and ITC data suggest that A12 is able to bind both the SOS response control proteins with micromolar affinity.

### A12 inhibition on LexA_Pa_ requires disulfide-mediated covalent binding

Since A12, 3-(2-sulfanylanilino)propanoic acid, is characterized by the presence of a thiol group and both RecA_Pa_ and LexA_Pa_ have some cysteine residues not involved in intramolecular disulfide bridges, the possibility that A12 targets the SOS system via the formation of disulfides was investigated. Moreover, we considered that in the experimental conditions used throughout this work A12 could form a disulfide-mediated adduct, thus converting back to its synthesis intermediate 3-[2-[[2-(2-carboxyethylamino)phenyl]disulfanyl]anilino]propanoic acid, which might represent the biologically active form. These possibilities were investigated by mass spectrometry (MS) and fluorescence polarization assays (FP).

First, ESI-MS analysis of A12 (2 mM, diluted in the same buffer used for FP assays) revealed a predominant species with a molecular weight of 394.17 ± 0.05 Da (Fig. 3 A), which is compatible with that of 3-[2-[[2-(2-carboxyethylamino)phenyl]disulfanyl]anilino]propanoic acid (392.5 Da). The same analysis was conducted after the treatment of A12 with 2 mM DTT (Fig. 3 B). In this sample, the MS signals corresponding to the disulfide adduct were absent, leading to the conclusion that the dimerization of A12 by the formation of a disulfide bridge can occur under physiological conditions. When in monomeric form, the compound is likely degraded and/or not detectable by MS under the tested experimental conditions.

**Fig. 3:**
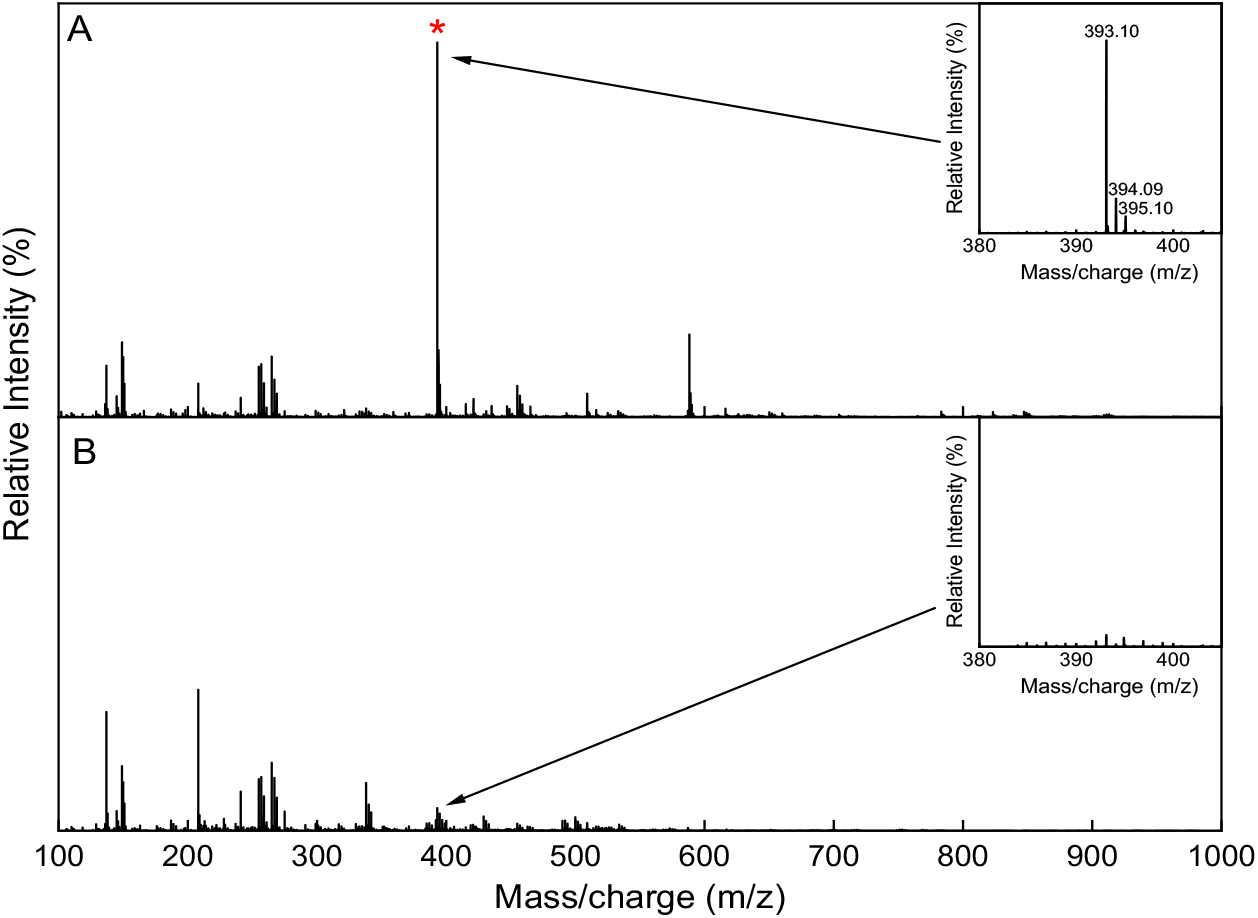
Positive-ion ESI mass spectrum of A12. (A) A12 in the absence of DTT showing dimeric form of the compound (red star). Inset: zoom of the region showing the charge distribution of the chemical compound. (B) A12 incubated with DTT (2 mM) showing the absence of the dimer in reducing condition. Insert: zoom of the region previously containing the dimeric signal

Then, ESI-MS analysis was carried out on both RecA_Pa_ and LexA_Pa_ alone and pre-incubated with A12 (referred to as RecA_Pa_-A12 and LexA_Pa_-A12, respectively).

Compared to the unreacted control (which displayed a single protein species; Fig. 4A and Fig. S2 A), the mass spectrum of RecA_Pa_-A12 showed the presence of two protein species (Fig. 4B and Fig. S2 A) with a 179 Da mass difference. This difference is compatible with a condensation reaction between A12 and RecA_Pa_, with the release of a molecule of water. A tryptic digestion of RecA_Pa_-A12 was submitted to LC-MS analysis (Fig. S2 A-B), revealing that the chemical modification locates on the 270-285 fragment of RecA_Pa_ (TGEIIDLGVQLGLVEK; residues 285-300 in the 6xHis-tagged protein).

**Fig. 4:**
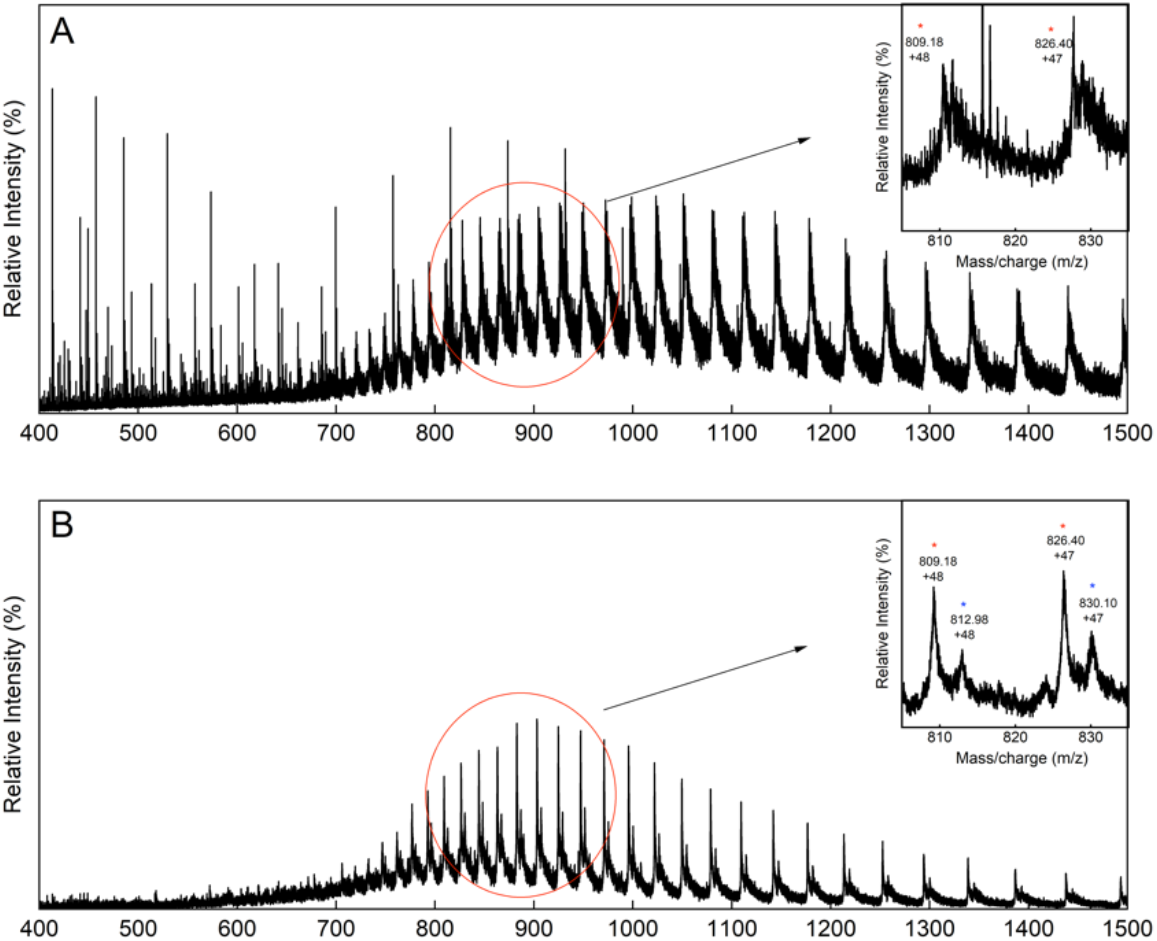
MS analysis of RecA_Pa_ incubated with A12. (A) Mass spectrum of RecA_Pa_. (B) Mass spectrum of RecA_Pa_-A12, showing the appearance of a second mass distribution. Insets: zoom of the range 800-900 m/z, showing the signals of the protein distribution, corresponding to RecA_Pa_ (red star) and to the A12-bound RecA_Pa_ (blue star), with a mass delta of 179 Da.

The mass spectrum of LexA_Pa_ (Fig. 5 A and Fig. S2 A) shows a single species corresponding to the unmodified protein. Conversely, MS analysis of LexA_Pa_-A12 revealed the presence of two protein species: one with the expected molecular weight for LexA_Pa_ (Fig. 5 B, red star; Fig. S2 A) and the other with a molecular weight 195 Da higher (Fig. 5 B, blue star). The latter probably corresponds to LexA_Pa_ covalently bound by A12 by a disulfide bond, as a 2 Da mass difference between A12 and the identified modification is compatible with thiols deprotonation upon disulfide formation.

**Fig. 5:**
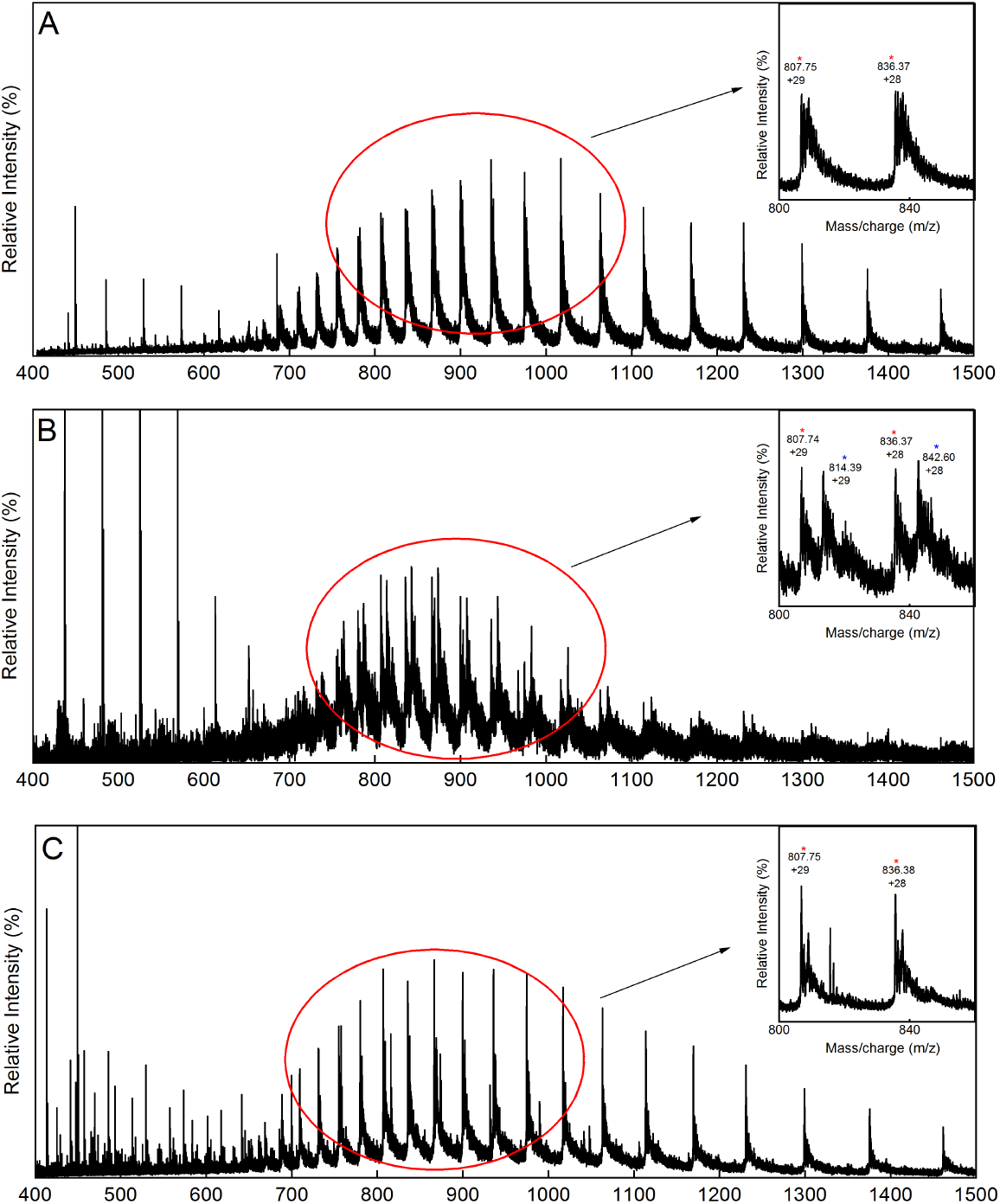
MS analysis of LexA_Pa_ incubated with A12. (A) Mass spectrum of LexA_Pa_. (B) Mass spectrum of LexA_Pa_-A12, showing the appearance of a second mass distribution. (C) Mass spectrum of LexA_Pa_-A12 treated with DTT, showing that the chemical modification is lost after the reducing treatment. Insets: zoom of the 800-900 m/z range, showing the signals of the protein distribution without (red star) and with (blu star) a delta mass of 195 Da.

A control analysis was performed in the presence of 5 mM DTT and the chemically modified LexA_Pa_ was absent (Fig. 5C and Fig. S2 A and S2 C), confirming that A12 covalent binding on LexA_Pa_ relies on an oxidation reaction.

Once again, tryptic digestion and comparative LC-MS analysis of LexA_Pa_ alone and co-incubated with A12 allowed the identification of the chemically modified peptide (Fig. S2B), corresponding to the 88-105 fragment (VAAGAPILAEQNIEES**C**R; residues 95-112 considering the 6xHisTag), which includes Cys104 (Cys111 in the 6xHis-tagged protein). The un-modified peptide has a molecular mass of 1869.93 ± 0.08 Da, while the modified one showed an increase in mass of 195 Da. These data led to the conclusion that A12 is able to bind LexA_Pa_ by the formation of disulfide bond at the level of Cys104.

We further investigated the relevance of disulfide-mediated chemical binding of A12 to LexA_Pa_ by the FP-based autoproteolysis assay under reducing conditions (2 mM TCEP). The reducing environment significantly diminished A12 inhibitory activity compared to a non-reducing condition (Fig. 6 A), confirming that redox equilibria have an impact on its mechanism of action.

**Fig. 6:**
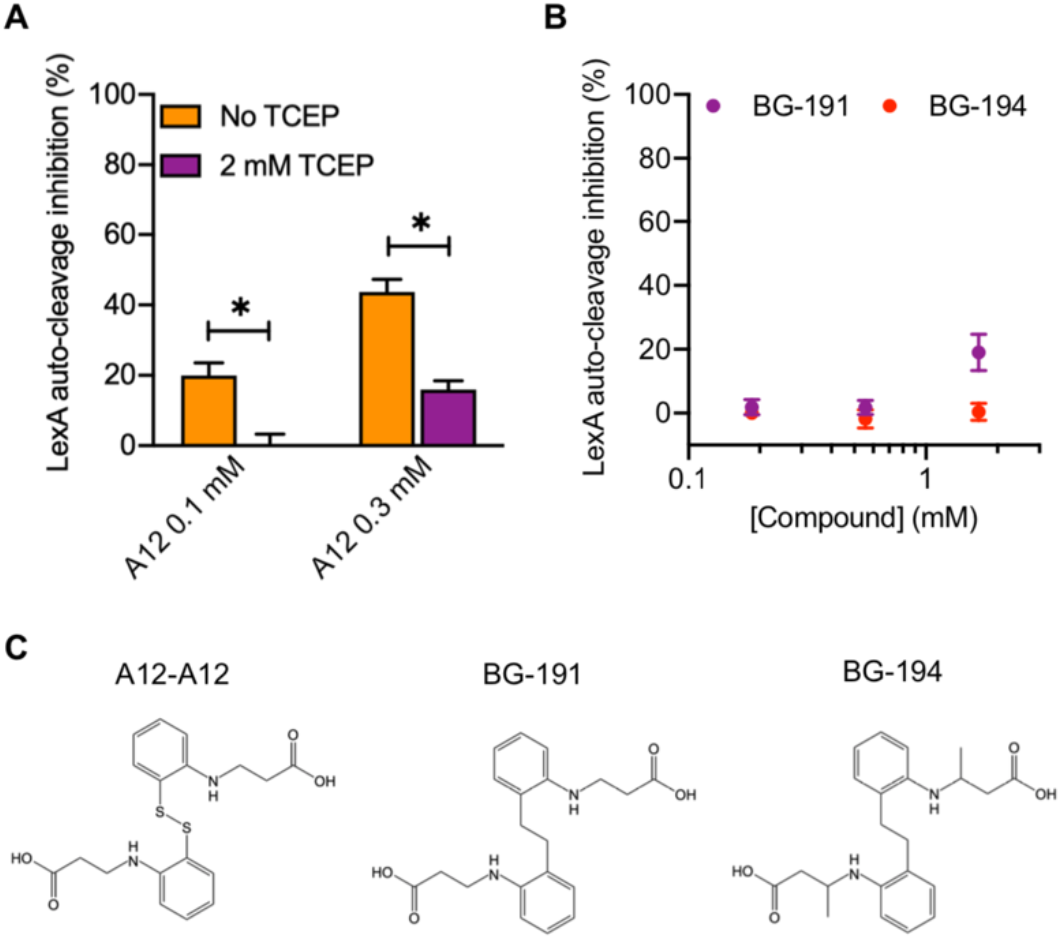
Activity testing of A12 in a reducing environment and of A12 derivatives. (A) LexA_Pa_ auto-cleavage percent inhibition obtained by the FP-based assay with different concentrations of A12, either in the presence or absence of a reducing agent (TCEP). (B) LexA_Pa_ auto-cleavage percent inhibition obtained by the FP-based LexA_Pa_ autoproteolysis assay with different concentrations of A12 derivatives. (C) Chemical structures of A12 covalent adduct and derivatives. Data represent the mean ± S.E.M. of three replicates. *P-value < 0.03.

To further exclude that the biologically active form of A12 is its covalent dimeric adduct, an analog corresponding to two A12 moieties devoid of sulfur atoms and linked by an ethylene arm (3,3’-((ethane-1,2-diylbis(2,1-phenylene))bis(azanediyl))dipropionic acid, referred to as BG-191) was synthesized and examined by the same FP-based LexA_Pa_ self-cleavage assay. A slightly modified congener (3,3’-((ethane-1,2-diylbis(2,1-phenylene))bis(azanediyl))dibutyric acid, referred to as BG-194) was tested as well. Both compounds displayed a severely lowered activity compared to A12 in a range of concentrations between 0.2 and 2 mM. While BG-194 did not display any inhibitory activity on LexA_Pa_, BG-191 reached around 20% inhibition at the highest concentration tested (Fig. 6 B). Conversely, as discussed above, A12 almost completely prevented LexA_Pa_ autoproteolysis at 1 mM. Taken together, these observations suggest that the observed activity of A12 relies on the direct binding of its monomeric form to LexA_Pa_ via disulfide bridging (Fig. 7 A-B).

**Fig. 7:**
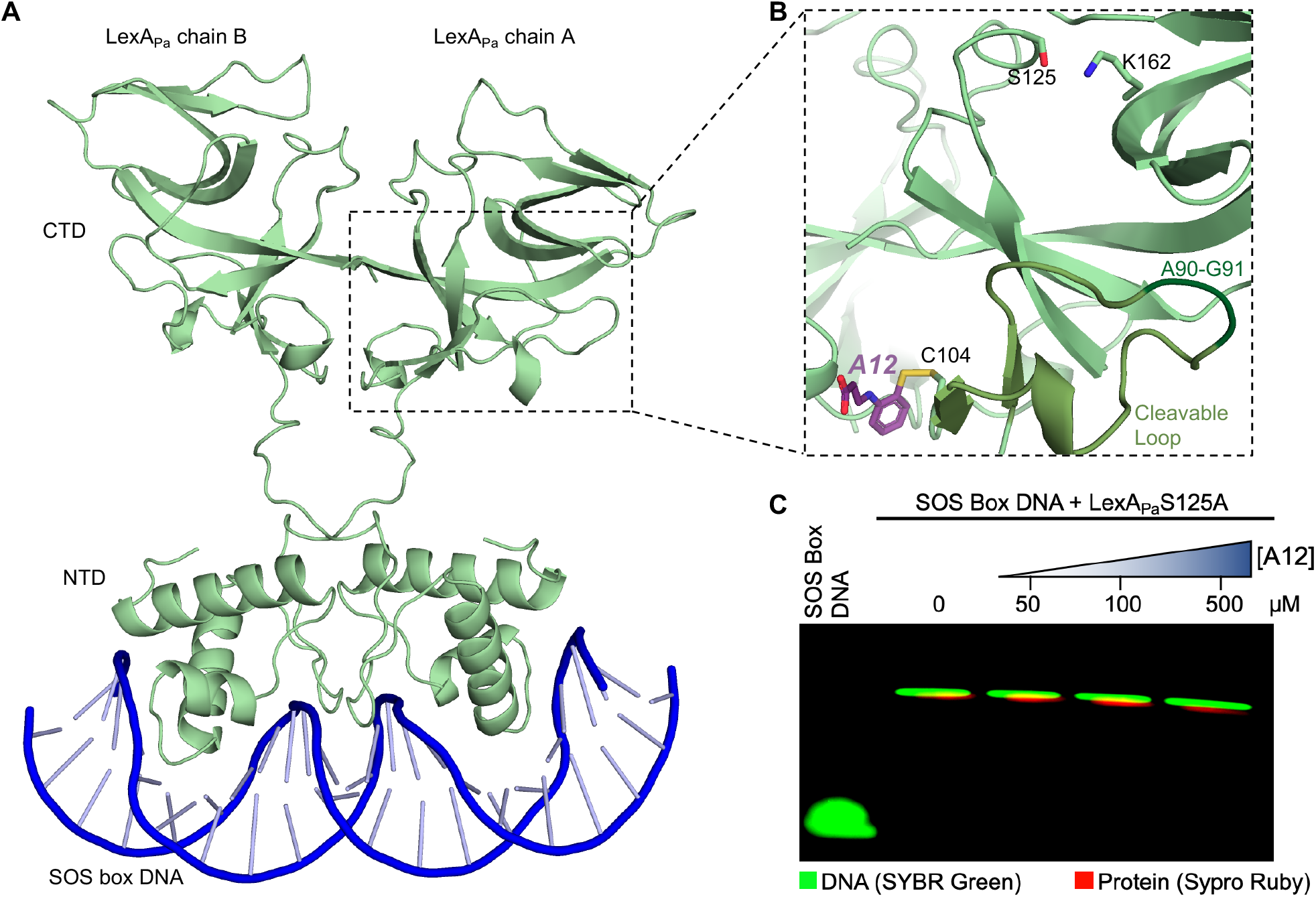
A12 binding site on LexA_Pa_. (A) AlphaFold3 model of full-length LexA_Pa_ bound to SOS box dsDNA. (B) Zoom on A12 binding site on LexA_Pa_^CTD^ (light green; PDB: 8B0V) at the base of the cleavable loop (medium green; the cleavage site A90-G91 is depicted in dark green) and close to the dimerization interface. The catalytic dyad S125/K162 is shown as sticks. (C) EMSA showing LexA_Pa_^S125A^ binding to SOS box dsDNA, in the presence of increasing concentrations of A12.

**Fig. 7:**
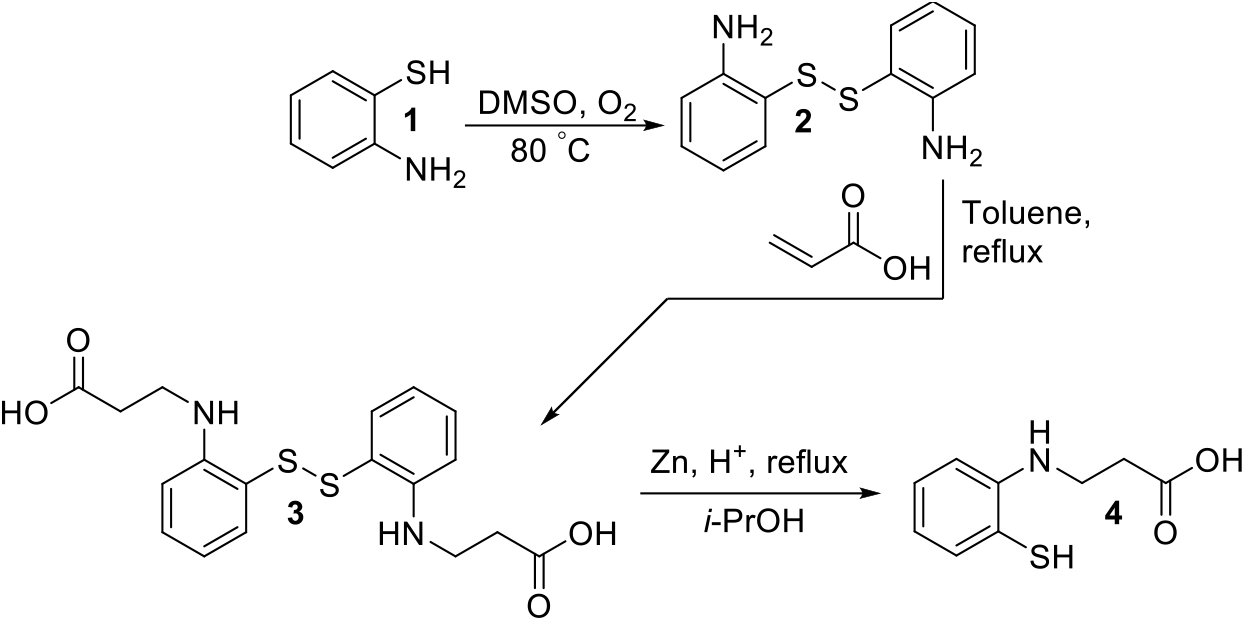
Chemical synthesis of A12 (compound 4)

### A12 does not impair LexA_Pa_ DNA-binding activity

LexA_Pa_ DNA-binding activity – which has to be preserved for an anti-SOS strategy to be effective – resides on the NTD but requires dimerization via the CTD. In this context, to rule out any interfering effect of A12 on LexA_Pa_ binding to SOS-box DNA, an electrophoretic mobility shift assay (EMSA) was performed by incubating LexA_Pa_ with operator dsDNA sequences and different doses of A12, before analyzing the mixtures on native polyacrylamide gels. Even in the presence of a 60-fold molar excess of A12, LexA_Pa_ preserved an unchanged ability to bind SOS-box dsDNA, as confirmed by the conserved band shifts to a higher molecular weight (Fig. 7 C).

## Discussion

In the present work, potential inhibitors of the *P. aeruginosa* SOS system have been searched among small molecules by a medium-throughput screening *in vitro*. One hit inhibitor of RecA_Pa_*-induced LexA_Pa_ autoproteolysis was discovered and two different assays – relying on FP and SDS-PAGE to follow LexA_Pa_ self-cleavage – were used to characterize its dose-response relationship. Obtained IC_50_ values lies in the mid micromolar range, with differences in estimates attributed to intrinsic dissimilarities of the two techniques. In particular, different LexA_Pa_ variants (FlAsH-LexA_Pa_^CTD^ *vs* full-length LexA_Pa_) were used in the two assays, which might account for diverse affinities for A12. In the FP-based assay, the presence of thiol-reactive species, such as unreacted FlAsH-EDT_2_ and unlabeled 4Cys-tagged LexA_Pa_^CTD^, could account for partial A12 depletion and interference with the system. These species may form undesired covalent bonds with the A12 thiol group, likely competing with its inhibitory reaction on FlAsH-LexA_Pa_ and potentially leading to underestimation of inhibitory activity.

The hit-to-lead optimization of A12 by targeted pharmacophore exploration would benefit from insights on A12 mechanism of action. To this aim, A12 interaction with RecA_Pa_ and LexA_Pa_ underwent biochemical and biophysical exploration by means of differential scanning fluorimetry (DSF) and isothermal titration calorimetry (ITC). The former technique revealed that A12 induced T_m_ variations on both the tested proteins, with a particular stabilizing effect on RecA_Pa_, a clear indication of binding [10]. Conversely, increasing concentrations of A12 transformed the biphasic melting curve of LexA_Pa_ in a monophasic curve with a generally higher sensitivity to thermal denaturation. Biphasic melting curves are typical of proteins that either consist of two independently folding domains [10] or assume a dimeric organization [11]. Both cases could potentially apply to LexA_Pa_, as it is composed of two domains connected by a long and flexible linker and it behaves as a dimer in solution [4]. Therefore, DSF data suggest that A12 could bind LexA_Pa_ in a region at the boundary of the C-terminal and N-terminal domains or close to the dimerization surface. On the other hand, ITC confirmed that A12 binds to both LexA_Pa_ and RecA_Pa_, with approximately 2 equivalent binding sites on each protein, but a higher affinity toward the transcriptional repressor LexA_Pa_ (*KD*^*App*^ of 7.4 µM) than for the co-activator RecA_Pa_ (*KD*^*App*^ of 18 µM).

A12 underwent mass spectrometry (MS) analysis, showing that it mainly exists as a disulfide bridged self-dimer in solution, reverting back to its synthesis intermediate. More notably, MS analysis evidenced that A12 binds RecA_Pa_ and LexA_Pa_ covalently, revealing a different binding mechanism on the two proteins: a condensation reaction in the former case and a disulfide bridging in the latter one. Further activity testing *in vitro* in the presence of a reducing agent confirmed that the formation of disulfides with LexA_Pa_ is crucial for A12 activity. To confirm that the species mainly responsible for LexA binding and inhibition is the reduced A12 monomer, a molecule mimicking an A12 self-dimer devoid of thiols was tested and resulted inactive in a broad range of concentrations, excluding that the biologically active form could be represented by the dimeric A12 adduct. In light of such results, we hypothesize that the oxidized A12 dimer and its monomeric reduced form define an equilibrium in solution that limits the concentration of monomeric A12 available to bind and inhibit LexA_Pa_. This suggests that the affinity of A12 toward LexA_Pa_ might be higher than the calculated values and further explains the apparent 2-fold molar ratio required to reach half saturation in ITC.

MS analysis allowed to identify Cys104 as the A12 binding site on LexA_Pa_: such residue is located in the C-terminal autoproteolytic domain, at the basis of the cleavable loop and in close proximity to LexA_Pa_ dimerization interface, as observable in the previously solved structure of LexA_Pa_ (PDB 8B0V) [4]. This allowed us to suggest that A12’s mechanism of action could rely on the constraint of LexA_Pa_ cleavable loop in an inactive conformation or the alteration of LexA_Pa_ dimeric architecture. To rule out the possibility that A12 might destabilize the conformation of LexA_Pa_ required for binding to *SOS box* dsDNA, an electrophoretic mobility shift assay was performed. The obtained results show that A12 treatment maintains LexA_Pa_ DNA-binding activity. Besides validating A12 as a hit SOS inhibitor, these data revealed that Cys104 can be targeted as a site for blocking LexA_Pa_ autocleavage while keeping the SOS pathway silenced.

In conclusion, the screening and validation campaign described here led to the identification of a new potential LexA_Pa_ targetable site and a molecular scaffold for SOS pharmacological suppression. Our findings open the way for further investigation and optimization of A12, to derive a potent and specific lead for future applications against the spreading of multi-resistant *P. aeruginosa* infections.

## Materials and methods

### Gene cloning and mutagenesis

All the plasmid vectors described in this work have been obtained as previously described [4]. Briefly, the genomic DNA of *P. aeruginosa* ATCC 27853 was purified from an overnight liquid culture and used as the template to PCR-amplify the coding sequences of RecA_Pa_ (UniProtKB: P08280) and LexA_Pa_ (UniProtKB: P37452), using primer couples RecA_Pa_pColi.For/Rev and LexA_Pa.For/Rev (Supplementary Table 1). The obtained amplicons were cloned in pColiExpressI (Canvax) and pETite C-His Kan vector (Lucigen), respectively, according to manufacturers’ instructions, obtaining the plasmids pColiXP-RecA_Pa_ and pETite-LexA_Pa_.

The QuikChange Site-Directed Mutagenesis Kit protocol (Agilent Technologies) was applied to pETite-LexA_Pa_, using the mutagenic primers LexA_Pa_S125A.For and LexA_Pa_S125A.Rev (Supplementary Table 1), obtaining the pETite-LexA_Pa_S125A plasmid vector.

The coding sequence of TetraCys-tagged LexA_Pa_ C-terminal domain (CTD) was amplified using primers LexA_Pa_CTD_4Cys.For and LexA_Pa_CTD_4Cys.Rev (Supplementary Table 1) and cloned in pETite N-His SUMO Kan Vector (Lucigen) according to manufacturer’s instructions, obtaining the plasmid pETite-SUMO-4Cys-LexA_Pa_^CTD^.

### Recombinant proteins expression and purification

All the proteins (6His-RecA_Pa_, 6His-LexA_Pa_ wt and S125A mutant and FlAsH-LexA_Pa_^CTD^) were recombinantly expressed in *E. coli* BL21(DE3) cells, transformed with the appropriate plasmid vector, as previously reported [4]. Briefly, 2-liter cultures in LB broth were set up. Protein overexpression was induced by adding 1 mM IPTG to bacterial cultures in the late exponential growth phase (OD_600_ 0.6-0.8) and was carried out overnight at room temperature under vigorous shaking (180 rpm).

#### RecA_Pa_

Cells were harvested by centrifugation and resuspended in RecA_Pa_ Buffer A (10 mM Hepes, 300 mM NaCl, 10% v/v Glycerol, 20 mM Imidazole, pH 8.0) supplemented with Protease Inhibitors Cocktail (SERVA). Bacterial cells lysis was performed by sonication. Cell debris were removed by centrifugation and the lysate soluble fraction was loaded on a 5 mL HisTrap Excel IMAC column (Cytiva). After extensively washing the column with RecA_Pa_ Buffer A and with 50 mM imidazole in Buffer A, His-tagged RecA_Pa_ was eluted by linearly raising the imidazole concentration in the eluent from 50 mM to 500 mM in 3 column volumes.

IMAC fractions showing RecA_Pa_ as the main protein component in SDS-PAGE analysis were pooled together, concentrated using a Vivaspin Turbo Ultrafiltration unit (10 kDa MWCO; Sartorius) and buffer-exchanged to RecA_Pa_ Storage Buffer (10 mM Hepes, 300 mM NaCl, 10% Glycerol, 1 mM MgCl_2_, 1 mM DTT, pH 7.0) by a HiTrap Desalting column (Cytiva) before storage at -80 °C for future usage in *in vitro* assays.

#### LexA_Pa_ wild-type and S125A uncleavable mutant

The same protocol was used for recombinant expression and purification of wild-type and mutant LexA_Pa_ variants.

Cells were harvested by centrifugation and resuspended in LexA_Pa_ Lysis Buffer (20 mM Tris-HCl, 150 mM NaCl, 20 mM Imidazole, 10 % v/v Glycerol, pH 7.5) supplemented with Protease Inhibitors Cocktail (SERVA), 500 U of benzonase nuclease (Merck) and 1.5 mM MgCl_2_. Bacterial cells lysis was performed by sonication and the crude lysate was incubated 30 minutes at 4 °C to allow benzonase-mediated DNA digestion. Cell debris were removed by centrifugation and the lysate soluble fraction was loaded on a 1 mL HisTrap Excel IMAC column (Cytiva). After extensively washing the column with LexA_Pa_ Buffer A (20 mM Tris-HCl, 150 mM NaCl, 10 % v/v Glycerol, pH 7.5) and with 20 mM imidazole in Buffer A, His-tagged LexA_Pa_ was eluted by linearly raising the imidazole concentration in the eluent from 20 mM to 500 mM in 10 column volumes.

IMAC fractions showing LexA_Pa_ as the main protein component by SDS-PAGE analysis were pooled together, concentrated using a Vivaspin Turbo Ultrafiltration unit (5 kDa MWCO; Sartorius) and buffer-exchanged to LexA_Pa_ Buffer A by a HiPrep 26/10 desalting column (Cytiva). 6His-LexA_Pa_ variants were stored at -80 °C for future usage in *in vitro* assays.

#### FlAsH-LexA_Pa_^CTD^

Cells were harvested by centrifugation and resuspended in FlAsH-LexA_Pa_ Lysis Buffer (20 mM Tris-HCl, 150 mM NaCl, 20 mM Imidazole, 10 % v/v Glycerol, 0.1 mM DTT, pH 7.5) supplemented with Protease Inhibitors Cocktail (SERVA). Bacterial cells lysis was performed by sonication.

Cell debris were removed by centrifugation and the lysate soluble fraction was loaded on a 1 mL HisTrap Excel IMAC column (Cytiva). After extensively washing the column with FlAsH-LexA_Pa_^CTD^ Buffer A (20 mM Tris-HCl, 150 mM NaCl, 10 % v/v Glycerol, 0.1 mM DTT, pH 7.5) and with 20 mM imidazole in Buffer A, 6His-SUMO-TetraCys-LexA_Pa_^CTD^ was eluted by linearly raising the imidazole concentration in the eluent from 20 mM to 500 mM in 10 column volumes.

IMAC fractions showing 6His-SUMO-TetraCys-LexA_Pa_^CTD^ as the main protein component by SDS-PAGE analysis were pooled together, diluted three times in FlAsH-LexA_Pa_^CTD^ Buffer A (to reduce imidazole concentration below 150 mM) and supplemented by 1 mM DTT, 1 mM EDTA, 0.1% v/v NP-40, and an excess of Expresso Sumo Protease. Following a 2-hours incubation at room temperature with gentle shaking, 100 μM FlAsH-EDT_2_ was added to the reaction mix and the incubation was prolonged overnight at 4 °C. The mixture was first concentrated using a Vivaspin Turbo Ultrafiltration unit (5 kDa MWCO; Sartorius) and buffer-exchanged to FlAsH-LexA_Pa_^CTD^ Final Buffer (20 mM tris-HCl, 150 mM NaCl, 10% v/v Glycerol, pH 7.5) by a PD-10 desalting column (Cytiva). Then, to remove 6His-SUMO fragments and uncleaved protein constructs from the final sample, the mixture was passed through a 1 mL HisTrap Excel IMAC column (Cytiva) and the flowthrough was recovered. FlAsH-LexA_Pa_^CTD^ was stored at -80 °C for future usage in *in vitro* assays.

### RecA_Pa_ activation

Biologically active RecA_Pa_/ssDNA/ATPγS nucleoprotein filaments (RecA_Pa_*) were produced by incubating 6His-RecA_Pa_ with SKBT25-18mer ssDNA ([RecA_Pa_]:[18mer]=3.5:1) and 1 mM ATP S overnight at 4 °C.

### FP-based library screening and hits validation

A custom-made library of 318 chemical fragments (25 mg/mL stock solutions in DMSO), provided by P. Kavaliauskas (Weill Cornell Medicine of Cornell University) [6] was screened *in vitro* for their ability to inhibit RecA_Pa_*-mediated LexA_Pa_ autoproteolysis by a Fluorescence Polarization (FP)-based assay [5,7,8]. 20 μL reactions were set up in Nunc 384-Well Black microplates and final reaction mixtures included 1 μM FlAsH-LexA_Pa_^CTD^, the compounds (5% DMSO in all the samples, including the controls devoid of compounds) and 1 μM RecA_Pa_* (FP Buffer in the negative control; 30 mM Hepes, 150 mM NaCl, pH 7.1). The initial screening was performed on two dilutions of the compounds (25 and 125 μg/mL). The FP signal was measured after a 1-hour incubation at 37 °C by an EnVision MultiMode Plate Reader (Perkin Elmer) equipped with opportune FP filters and mirrors. To obtain LexA_pa_^CTD^ autoproteolysis percent inhibition, FP data were scaled from 0% (corresponding to the positive control sample) to 100% (negative control).

Compounds displaying an inhibition of at least 50% at the highest concentration and a rather consistent increase in activity at the two tested concentrations were selected as the hits.

The same FP assay was carried out on serial dilutions of selected compounds to determine a dose-response relationship. To evaluate the effect of a reducing environment, 2 mM Tris(2-carboxyethyl)phosphine (TCEP) was added to the reaction mixtures. The FP signal of FlAsH-LexA_pa_^CTD^ alone was detected for 5 min before adding the compounds. The system was monitored for additionally 5 min in the presence of the tested compounds, in order to observe potential aggregation events. Then, RecA_Pa_* was added and the autoproteolysis reaction was followed for 1 h at 37 °C (1 reading per minute). FlAsH-LexA_pa_^CTD^ autoproteolysis percent inhibition was calculated as mentioned above and plotted as a function of compounds concentration. Experimental data fitting was performed by GraphPad Prism 8 to obtain the absolute IC_50_.

### Chemical synthesis and characterization of compounds

#### General procedures

Reagents and solvents were purchased from Sigma-Aldrich and used without further purification. The reaction course and purity of the synthesized compounds were monitored by TLC using aluminium plates pre-coated with Silica gel with F254 nm (Merck KGaA, Germany). Melting points were determined with a B-540 melting point analyzer (Büchi Corporation, USA). IR spectra (ν, cm^-1^) were recorded on a Perkin–Elmer Spectrum BX FT–IR spectrometer (Perkin–Elmer Inc., MA, USA) using KBr pellets. NMR spectra were recorded on a Bruker Avance III (400, 101 MHz) spectrometer (Bruker BioSpin AG, Switzerland). Chemical shifts were reported in (δ) ppm relative to tetramethylsilane (TMS) with the residual solvent as internal reference ([D_6_]DMSO, δ = 2.50 ppm for ^1^H and δ = 39.5 ppm for ^13^C). NMR data are reported as follows: chemical shift, multiplicity, coupling constant [Hz], integration and assignment. Mass spectra were recorded on Waters SQ Detector 2 Spectrometer with an electrospray ionization (ESI) source. Elemental analyses (C, H, N) were conducted using the Elemental Analyzer CE-440 and results were in good agreement (±0.3%) with the calculated values.

#### Synthesis of A12

2-aminobenzenethiol was previously shown to react with α, β-unsaturated acids producing benzothiazepine derivatives upon heating [12,13]. 2-aminobenzenethiol (**1**) was oxidized in dimethyl sulfoxide with atmospheric oxygen at 80 ° C, producing 2-[(2-aminophenyl)disulfanyl]aniline (**2)**, which was then boiled for 13h at reflux with acrylic acid in toluene. The mixture was cooled down and the formed crystals were filtered off. 3-[2-[[2-(2-carboxyethylamino)phenyl]disulfanyl]anilino]propanoic acid (**3**) was purified by recrystallization from 2-propanol. A mixture of compound **3** (0.01 mol, 3.92 g), zinc dust (46 mmol, 3.00 g), 10% hydrochloric acid (20 ml) and 2-propanol (15 ml) was boiled for 15 minutes and filtered hot. Sodium acetate (0.5 g) was added to the mixture and the formed crystals were filtered off and purified by recrystallization from 2-propanol, obtaining 3-(2-sulfanylanilino)propanoic acid (compound **4**, referred to as *A12* throughout this paper; Fig. 7). Chemical characterization (^1^H-NMR, ^13^C-NMR, IR and ESI-MS spectra) of compounds **3** and **4** is reported in Fig. S3 and Fig. S4, respectively.

*3-[2-[[2-(2-carboxyethylamino)phenyl]disulfanyl]anilino]propanoic acid* (**3**); Fig. S3.

m. p. 117–118 °C.

^1^H NMR (400 MHz, DMSO-d_6_) δ: 2.45 (4H, t, *J*=6.8 Hz, 2NH<u>CH_2_</u>); 3.33 (4H, kv, *J*=6.6 Hz, 2CH_2_CO); 5.42 (2H, t, *J*=6.1 Hz, 2NH); 6.48 (2H, t, *J*=7.4 Hz, H_Ar_); 6.68 (2H, d, *J*=8.2 Hz, H_Ar_); 7.02 (2H, dd, *J*=7.6; 1.5 Hz, H_Ar_); 7.13-7.29 (2H, m, H_Ar_); 12.30 (2H, s, 2OH).

^13^C BMR (101 MHz, DMSO-d_6_) δ: 33.86 (2NH<u>CH_2_</u>); 39.17 (2CH_2_CO); 110.86, 116.43, 117.94, 132.42, 136.71, 149.19 (C_Ar_); 173.64 (2CO).

IR (KBr), ν(cm^-1^): 3382, 3050 (2OH, 2NH); 1702 (2CO).

Calcd. for C_18_H_20_N_2_O_4_S_2_, %: C 55.08; H 5.14; N 7.14. Found, %: C 55.05; H 5.03; N 7.03. C_18_H_20_N_2_O_4_S_2_[M^+^] exact mass = 392.09, MS (ESI) = 392.92.

*3-(2-sulfanylanilino)propanoic acid* (**4**); Fig. S4.

m. p. 132–133,5 °C.

^1^H NMR (400 MHz, DMSO-d_6_) δ: 2,46 (2H, t, *J*=6,3 Hz, NHC<u>H_2_</u>); 2,97-3,17 (2H, m, CH_2_CO); 4,35 (1H, s, SH); 5,51 (1H, br. s. , NH); 6,59-6,73 (1H, m, H_Ar_); 6,75–6,90 (2H, m, H_Ar_); 7,28 (1H, d, *J*=7,6 Hz, H_Ar_); 12,29 (1H, br. s., OH).

^13^C NMR (101 MHz, DMSO-d_6_) δ: 25,33 (NH<u>C</u>H_2_); 34,34 (<u>C</u>H_2_CO); 115,98; 117,44; 122,96; 132,79; 136,17; 145,19 (C_Ar_) ; 175,23 (CO).

IR (KBr), ν (cm^-1^): 3387 (OH); 3054 (NH); 1719 (CO).

Calcd. for C_9_H_11_NO_2_S, %: C 54,80; H 5,62; N 7,10. Found, %: C 54,75; H 5,57; N 7,07. C_9_H_11_NO_2_S[M^+^] exact mass = 197.05, MS (ESI) = 197.00.

#### Synthesis of BG-191 and BG-194

Dicarboxylic acids **6** and **7** were obtained by the reaction of 2,2’-(ethane-1,2-diyl)dianiline **(5)** with acrylic and crotonic acids, respectively (Fig. 8).

**Fig. 8:**
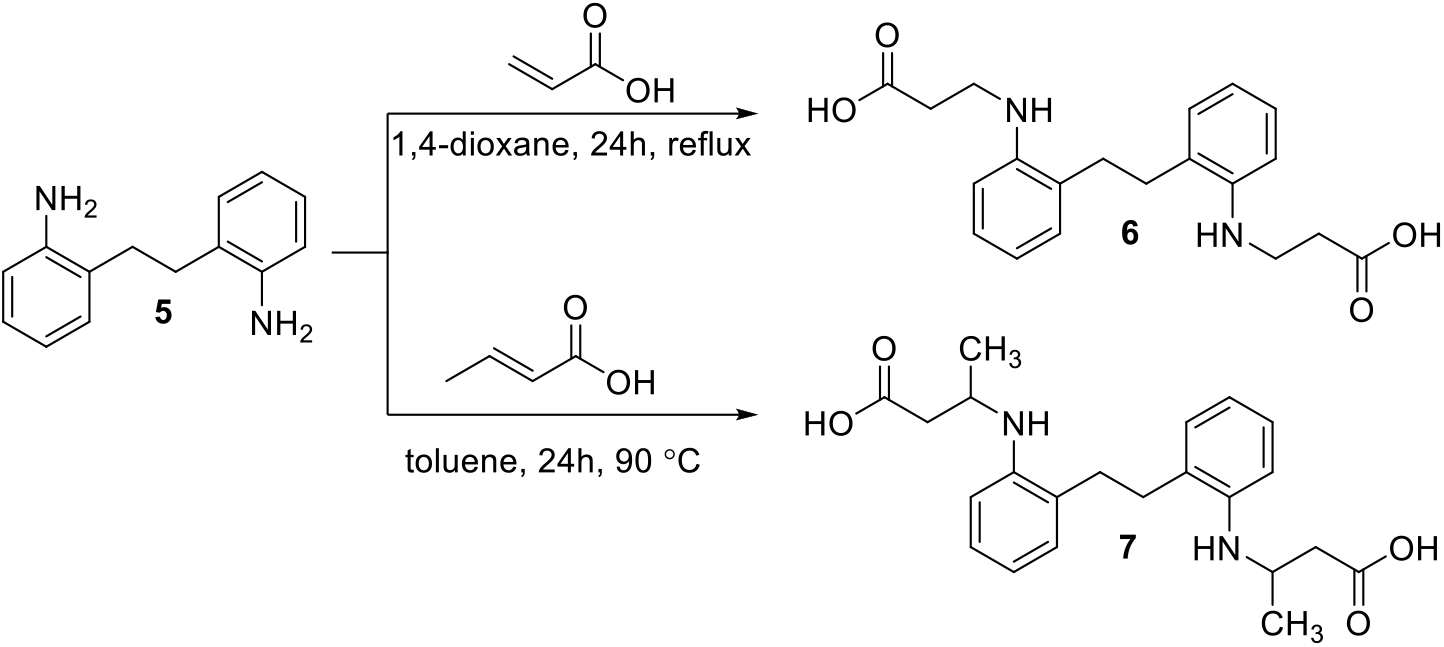
Chemical synthesis of BG-191 (compound 6) and BG-194 (compound 7).

A mixture of 2,2’-(ethane-1,2-diyl)dianiline **(5)** (4.7 mmol, 1 g), acrylic acid (9.8 mmol, 0.71 g) and 1,4-dioxane (5 mL) was heated at reflux for 24 h. The mixture was cooled down, diluted with diethyl ether and formed crystals were filtered off. Compound **6** was purified by recrystallization from 2-propanol and water.

A mixture of 2,2’-(ethane-1,2-diyl)dianiline **(5)** (4.7 mmol, 1 g), crotonic acid (14.1 mmol, 1.21 g) and toluene (15 mL) was heated at 90 °C for 24 h. The mixture was cooled down, 5% NaOH (20 mL) was added and extracted with diethyl ether. Aqueous solution was acidified with acetic acid to pH 6 and formed crystals were filtered off. Compound 7 was purified by recrystallization from 2-propanol. Chemical characterization (^1^H-NMR, ^13^C-NMR, IR and ESI-MS spectra) of compounds **6** and **7** is reported in Fig. S5 and Fig. S6, respectively.

*3,3’-((Ethane-1,2-diylbis(2,1-phenylene))bis(azanediyl))dipropionic acid* (**6**); Fig. S5.

m. p. 154–156 °C.

^1^H NMR (400 MHz, DMSO-d_6_) δ: 2.54 (4H, t, *J*=6.9 Hz, 2NH<u>CH_2_</u>); 2.66 (s, 4H, CH_2_CH_2_); 3,31 (4H, t, *J*=6.9 Hz, 2CH_2_CO); 4.94 (2H, br s, 2NH); 6.46−6.67 (4H, m, H_Ar_); 6.98−7.15 (4H, m, H_Ar_); 12.03 (2H, s, 2OH).

^13^C NMR (101 MHz, DMSO-d_6_) δ: 29.60 (2NH<u>CH_2_</u>); 33.63 (2CH_2_CO); 39.17 (CH_2_CH_2_); 109.65, 116.24, 126.10, 126.95, 128.84, 145.68 (C_Ar_); 173,74 (2CO).

IR (KBr), ν (cm^-1^): 3388, 3040 (2OH, 2NH); 1699 (2CO).

Calcd. for C_20_H_24_N_2_O_4_, %: C 67.40; H 6.79; N 7.86. Found, %: C 67.17; H 6.45; N 7.54. C_18_H_20_N_2_O_4_[M^+^] exact mass = 356.17, MS (ESI) = 356.91.

*3,3’-((Ethane-1,2-diylbis(2,1-phenylene))bis(azanediyl))dibutyric acid* (**7**); Fig. S6.

m. p. 106–108 °C.

^1^H NMR (400 MHz, DMSO-d_6_) δ: 1.18 (6H, d, *J*=6.3 Hz, 2CH_3_); 3.24−2.37 and 2.51−2.60 (4H, 2m, 2CH_2_CO) 2.66 (s, 4H, CH_2_CH_2_); 3.83 (2H, q, *J*=6.9 Hz, 2NH<u>CH_2_</u>); 5.54 (4H, br s, 2NH, 2OH); 6.45−6.61 (4H, m, H_Ar_); 6.94−7.10 (4H, m, H_Ar_).

^13^C NMR (101 MHz, DMSO-d_6_) δ: 20.00 (2CH_3_); 29.70, 29.98 (2NH<u>CH_2_</u>); 41.14 (2CH_2_CO); 45.18 (CH_2_CH_2_); 110.23, 114.79, 116.00, 116.43, 125.33, 126.30, 126.52, 126.85, 128.94, 129.12, 144.91, 146.11 (C_Ar_); 173,88 (2CO).

IR (KBr), ν (cm^-1^): 3370, 2965 (2OH, 2NH); 1690 (2CO).

Calcd. for C_22_H_28_N_2_O_4_, %: C 68.73; H 7.34; N 7.29. Found, %: C 68.52; H 7.21; N 7.11. C_22_H_28_N_2_O_4_[M^+^] exact mass = 384.20, MS (ESI) = 384.94.

### RecA_Pa_*-induced LexA_Pa_ autoproteolysis assay by SDS-PAGE

SDS-PAGE was used as an orthogonal technique to FP to validate the dose dependent inhibition of RecA_Pa_*-induced LexA_Pa_ autocleavage by the selected hit compound.

Full-length LexA_Pa_ was used instead of FlAsH-LexA_Pa_^CTD^, but reaction volumes, molar concentrations of reagents, incubation time and temperature were kept the same as in the FP-based assay. Following 1 h incubation at 37 °C in the presence of RecA_Pa_*, reactions were stopped by adding Laemmli solubilization buffer and freezing the samples at -20 °C.

Samples were boiled 5 minutes at 95 °C before being loaded on 4-12% polyacrylamide gels. After Coomassie staining of the gels, the intensity of the bands corresponding to LexA_Pa_ and its autoproteolytic fragments were quantified using ImageJ. Data obtained from LexA^CTD/NTD^ bands densitometry were used to estimate LexA self-cleavage percent inhibition (scaling all the data from 0%, positive control, to 100%, negative control). Autocleavage inhibition data were plotted as a function of compound concentration and submitted to nonlinear regression in GraphPad Prism 8 to determine the absolute IC_50_.

### Electrophoretic mobility shift assay

An electrophoretic mobility shift assay of the SOS-box dsDNA in the presence of full-length LexA_Pa_S125A and different concentrations of the hit compound was performed to observe potential interference effects of the selected drug on LexA_Pa_ binding to operator DNA sequences.

SOS-box dsDNA was produced by mixing equimolar amounts of AT-repeat_For and AT-repeat_Rev oligonucleotides (Supplementary Table 1) in annealing buffer (10 mM Tris-HCl, 50 mM NaCl, 1 mM EDTA, pH 7.5), heating at 95 °C for 5 minutes and then allowing the mixture to slowly cool down to room temperature overnight. The *E. coli* “AT-repeat” operator sequence [14] was used as it is fully compatible with the *P. aeruginosa* SOS-box consensus (CTG-N_2_-T-N_7_-CAG) [15].

1 μM SOS-box dsDNA was incubated either alone or with 8 μM 6His-LexA_Pa_S125A in EMSA buffer (50 mM Tris-HCl, 750 mM KCl, 0.5 mM DTT, 0.5 mM EDTA, pH 7.4). Different concentrations of the tested compound were added to the mixtures (0, 50, 100 or 500 μM; 10% v/v DMSO in all the reactions including controls) before a 40 minutes incubation at room temperature. Samples were treated with Purple DNA Loading Dye (New England Biolabs) and loaded on a NativePAGE 3-12% Polyacrylamide gel (Invitrogen). The gel was stained in SYBR Gold Nucleic Acid Stain (Invitrogen) for detecting DNA, then washed in deionized water and stained in SYPRO Ruby Protein Stain (Invitrogen) for detecting proteins. Obtained pictures were merged using ImageJ.

### Differential scanning fluorimetry

Differential Scanning Fluorimetry (DSF or “Thermofluor”) was performed on samples containing 4 μM RecA_Pa_ or LexA_Pa_S125A in FP Buffer (30 mM Hepes, 150 mM NaCl, pH 7.1), different concentrations of the selected compound and SyproOrange 8X (Life Technologies; a final concentration of 10% v/v DMSO was present in all the samples including controls).

Fluorescence of the SyproOrange dye was measured by a StepOne Real-Time PCR System (Applied Biosystems), while rising the temperature from 25 to 95 °C (0.5 °C/min). Normalized fluorescence intensity curves were compared to observe macroscopic differences in the recorded melting profiles, while derivative curves were analyzed to find exact protein melting temperatures (T_m_, corresponding to derivative curve minimum). Melting curves were repeated two times and duplicates were coherent if T_m_ differences were lower than 1 °C.

### Isothermal titration calorimetry

64 µM RecA_Pa_ and 58.5 µM LexA_Pa_S125A were titrated with the selected compound (2.5 mM solution) using a Microcal PEAQ-ITC instrument (Malvern Panalytical) at 25 °C with a stirring rate of 750 rpm. Proteins and ligand were dissolved in ITC Buffer (20 mM Tris-HCl, 150 mM NaCl, 5% v/v DMSO, pH 7.5). An initial 0.4 µL injection (excluded from data analysis) was followed by 24 injections of 1.5 µL with a spacing of 150 s between each addition. Blank experiments were performed injecting the titrant compound in protein-devoid ITC Buffer. Blank curves were subtracted to all the experiments. Data were analyzed using the MicroCal PEAQ-ITC Evaluation software (Malvern Panalytical). Integrated heat signals were fitted by a “one set of sites” binding model to obtain enthalpy changes (ΔH), dissociation constants (K_D_) and stoichiometry of binding.

### Mass spectrometry analysis

MS analysis was carried out on 5 µL samples of the compound A12 (2 mM), of the proteins 6His-LexA_Pa_ and 6His-RecA_Pa_ (50 µM), and of the same proteins incubated with 2 mM A12 (2h incubation at 25 °C, 10% v/v DMSO in 100 mM NaHCO_3_ pH 7.0). Aliquots of the latter samples were treated with 5 mM dithiothreitol (DTT) for 1h at 25 °C before being buffer-exchanged in 100 mM NaHCO_3_ pH 7.0. Samples were analyzed by an electrospray ionization (ESI) mass spectrometer equipped with a Xevo® G2-XS ESI-Q-TOF mass spectrometer (Waters Corporation, Milford, Massachusetts, USA). Measurements were carried out at 1.5–1.8 kV capillary voltage and 30–40 V cone voltage.

Tryptic digestion was obtained at a 1:20 trypsin to protein ratio (by weight) after the incubation of the proteins (6His-LexA_Pa_ and 6His-RecA_Pa_) with the chemical compound A12. The reaction was left at 37 °C overnight and then stopped by freezing at −20 °C. Proteolytic mixtures were analyzed by LC-MS, using an Agilent AdvanceBio Peptide Map column (2.1×150mm × 2.7 µm; Santa Clara, CA, USA) connected to an Acquity H-Class instrument (Waters Corporation, Milford, Massachusetts, USA). The elution was performed at a flow of 0.2 mL/min with the following acetonitrile/0.1% formic acid - water/0.1% formic acid gradient: 2–65%, 36 min, 65–98%, 2 min. Mass analyses were carried out with the same capillary voltage and cone voltage described before.

## Author Contributions

Conceptualization: F.V., D.T., P.K. and L.C.; Methodology: F.V., B.F., V.M., P.PdL., P.K., V.M., B.G., V.P. and L.C.; Investigation: F.V., B.F., V.M.; Formal Analysis: F.V., D.T., P.K. and L.C.; Writing – Original Draft: F.V., B.F. and L.C.; Writing – Review & Editing: F.V., P.PdL, D.T., V.M., B.G., V.P., P.K., A.P. and L.C.; Supervision: P.PdL., P.K., D.T. and L.C.; Resources: P.K., P.PdL., D.T., A.P. and L.C.; Project Administration: P.K., D.T. and L.C..

## Acknowledgments

The authors would like to thank Fondazione Cassa di Risparmio di Padova e Rovigo (Cariparo) and the Department of Biology of the University of Padova for supporting Filippo Vascon with a PhD scholarship and a postdoctoral fellowship.

## Supplementary Materials

**Table S1:**
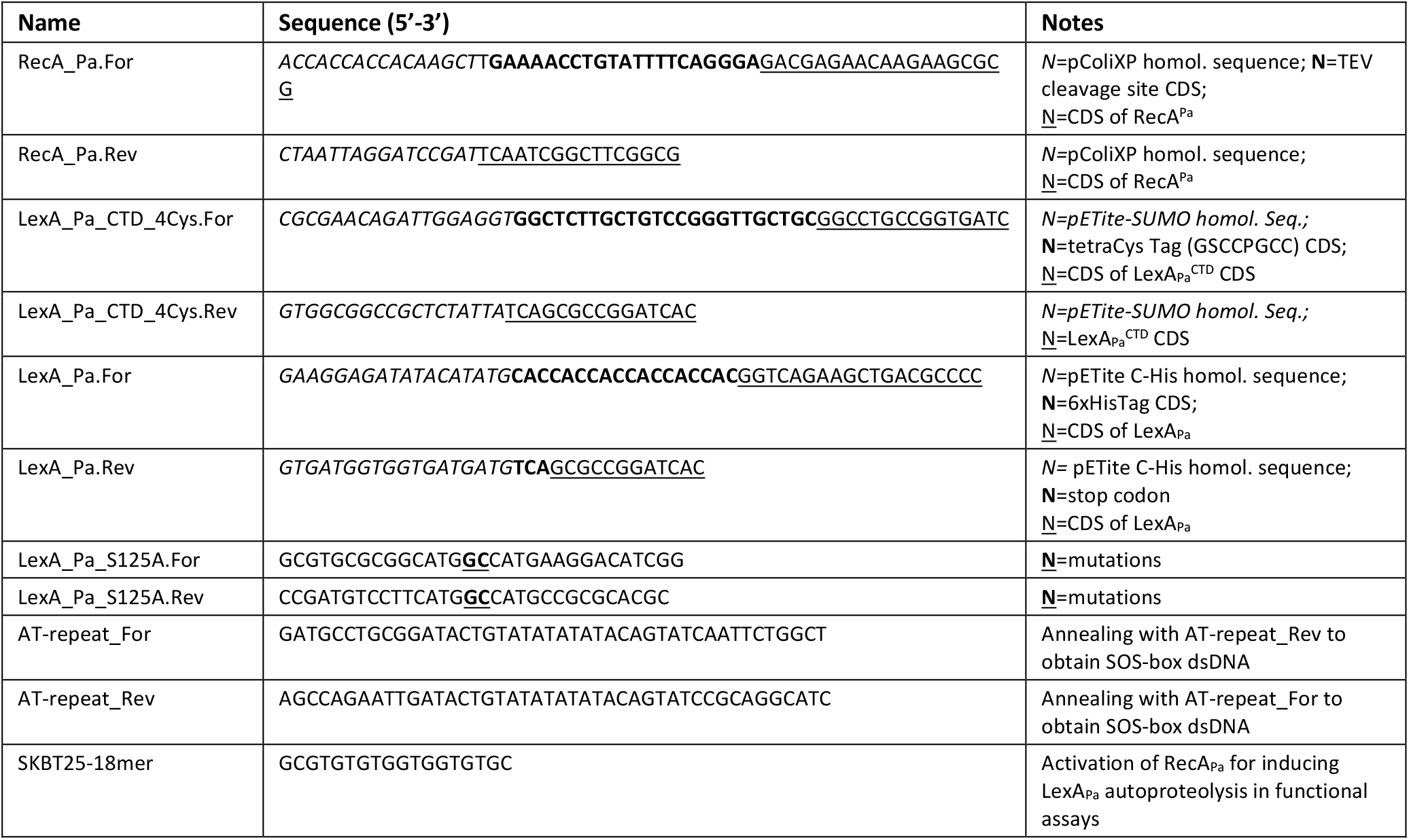
Oligonucleotides

**Fig. S1:**
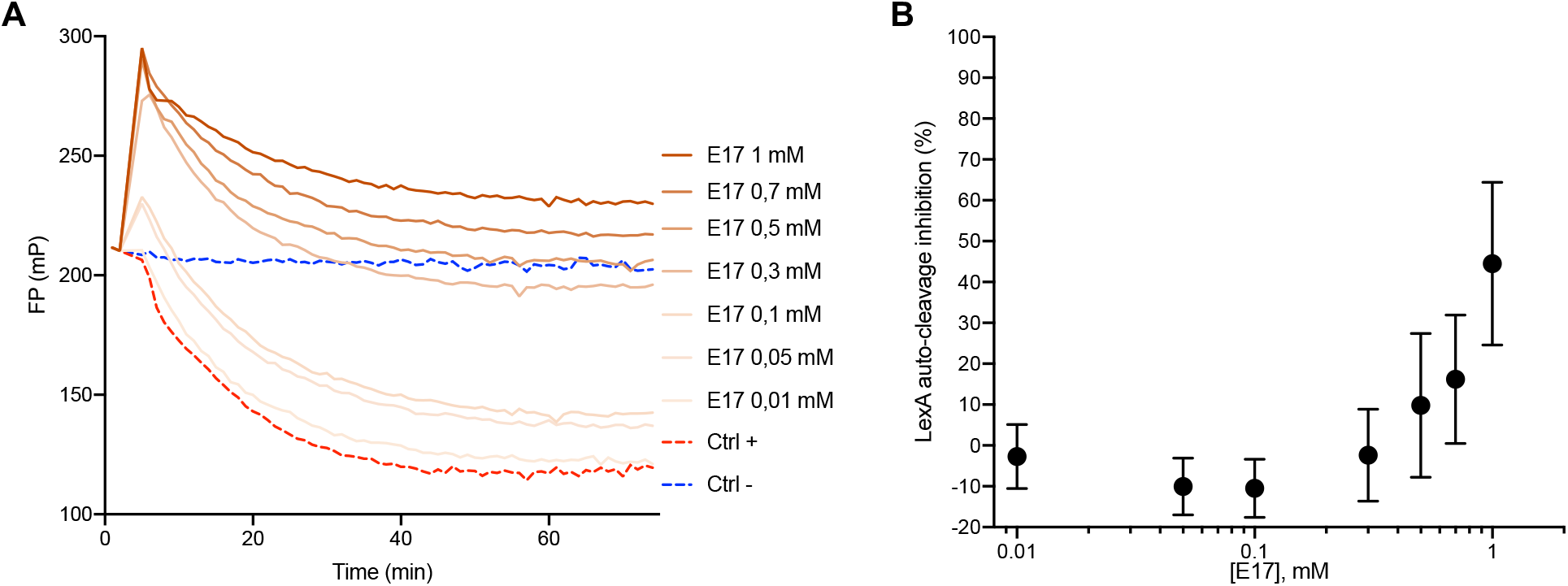
Characterization of compound E17 by the FP-based LexA autoproteolysis assay.

**Fig. S2:**
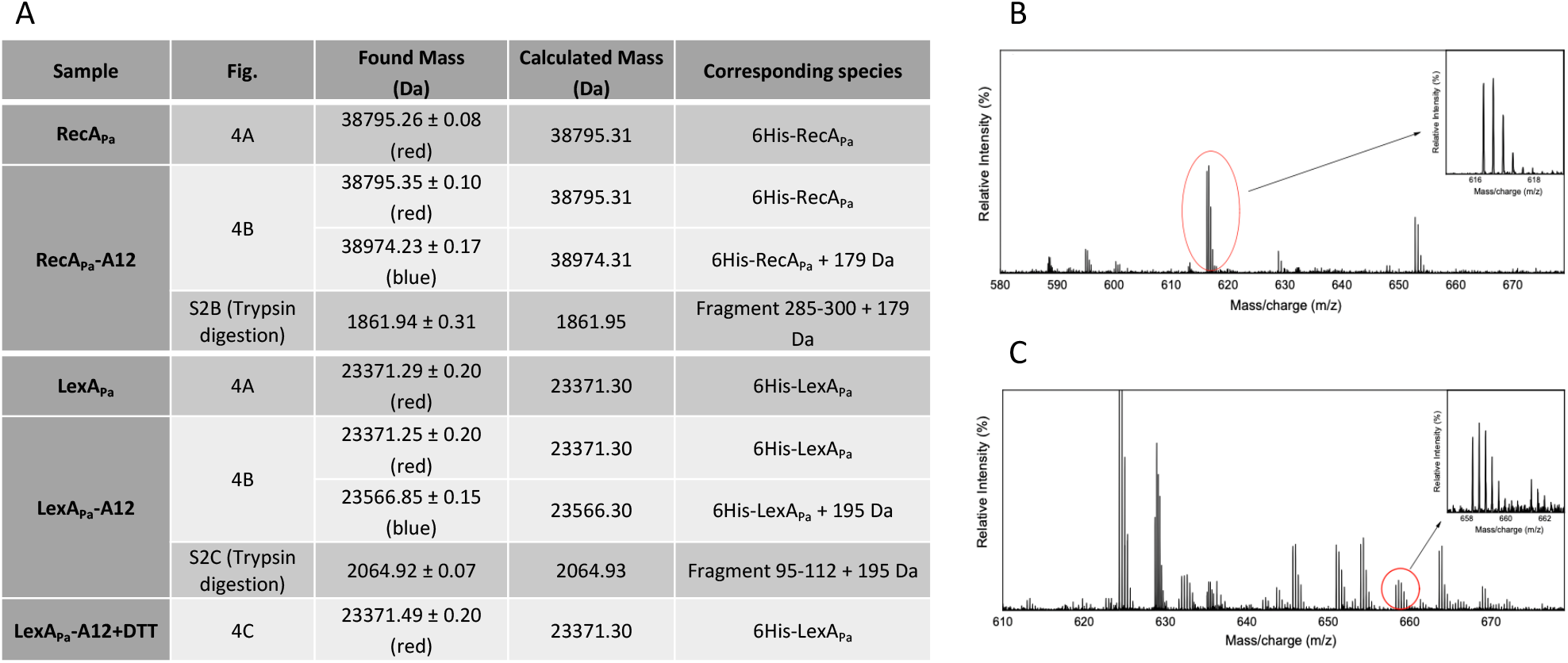
MS analysis of A12-modified RecA_Pa_ and LexA_Pa_. (A) Protein molecular masses found from MS analysis of the indicated samples. Molecular masses of found modified peptides in trypsin digestions are reported as well. Charge distributions of the chemically modified peptides of RecA_Pa_-A12 (B) and LexA_Pa_-A12 (C).

**Fig. S3:**
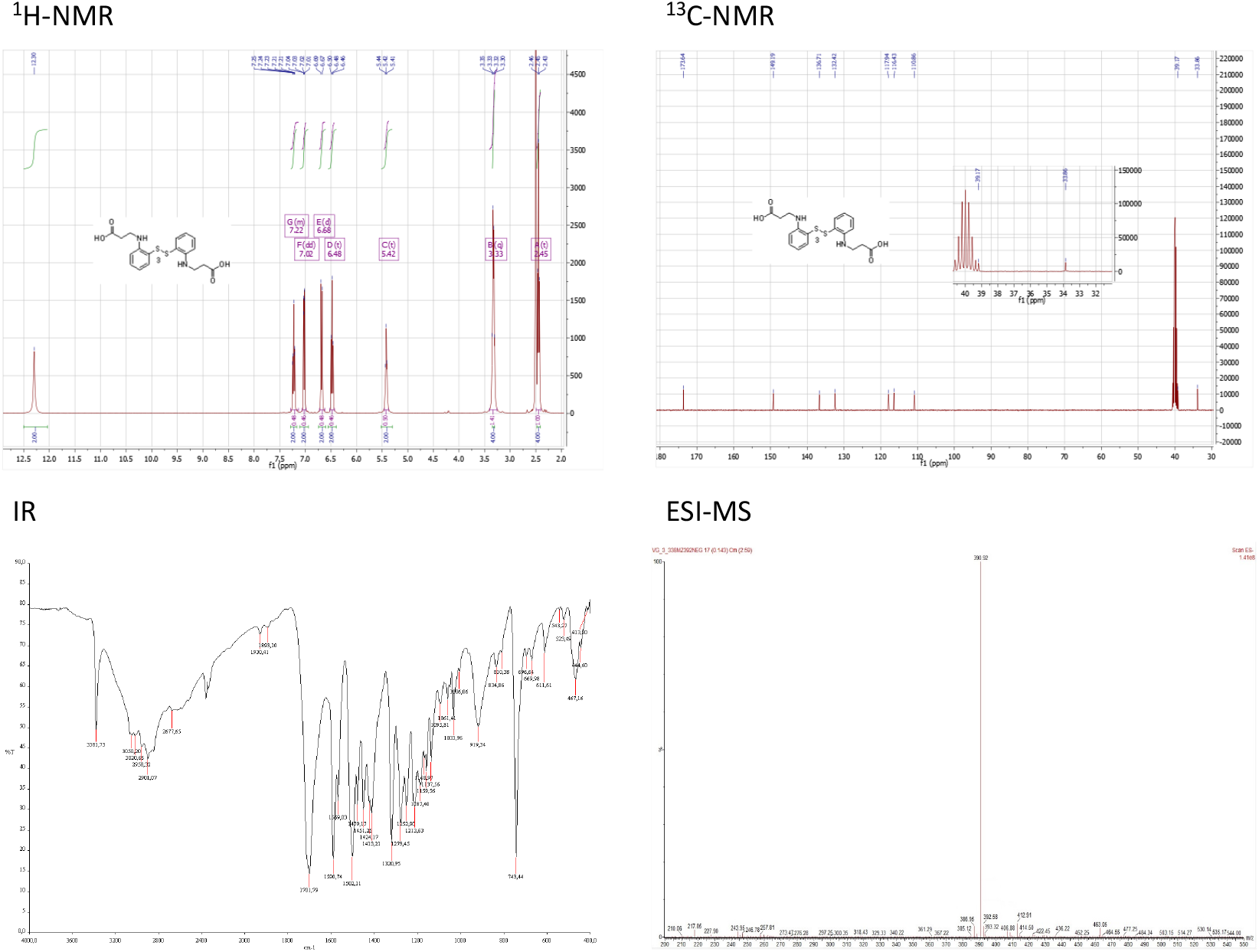
Characterization of compound 3 after chemical synthesis. (^1^H-NMR, ^13^C-NMR IR and ESI-MS spectra)

**Fig. S4:**
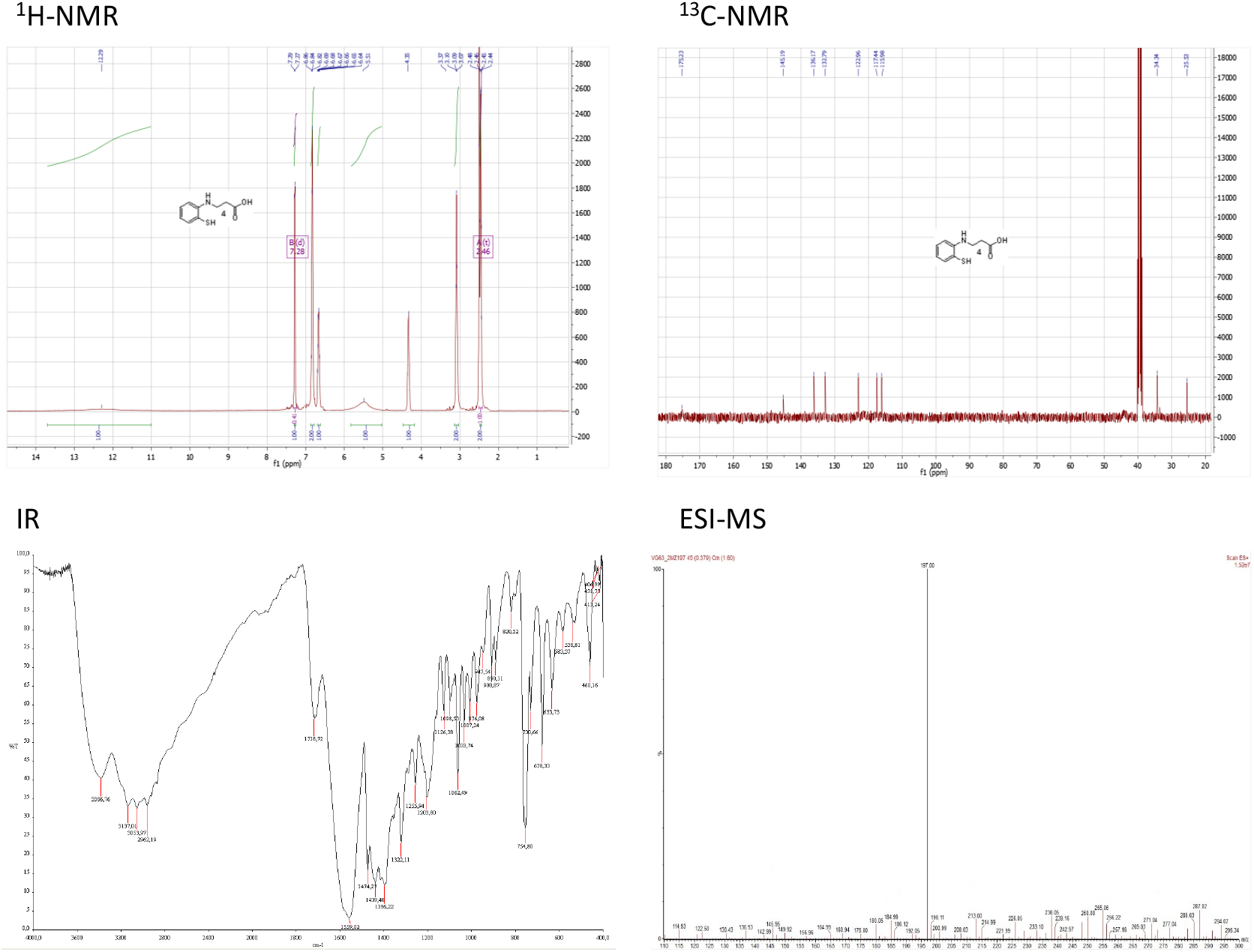
Characterization of compound 4 (A12) after chemical synthesis. (^1^H-NMR, ^13^C-NMR IR and ESI-MS spectra)

**Fig. S5:**
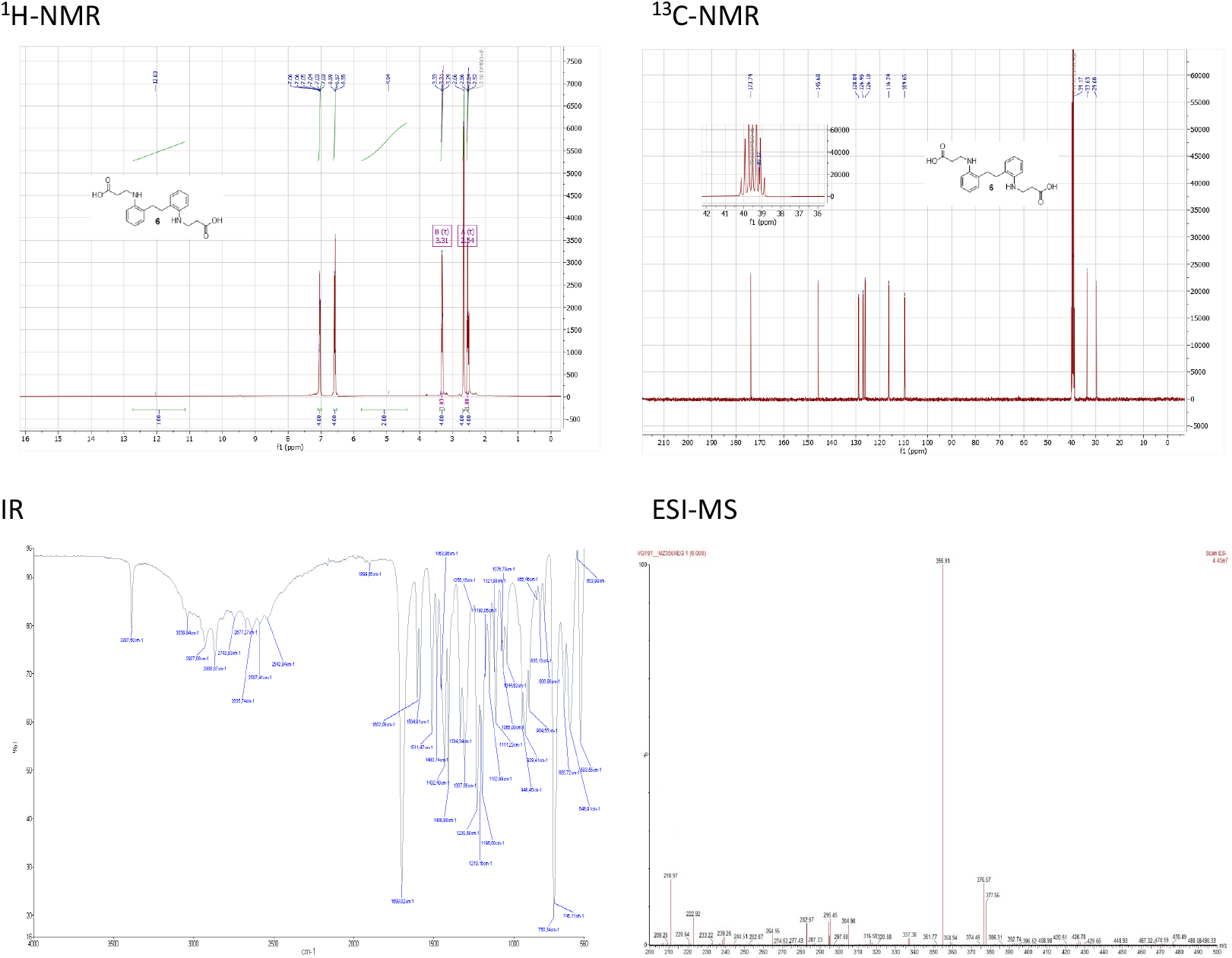
Characterization of compound 6 (BG-191) after chemical synthesis. (^1^H-NMR, ^13^C-NMR IR and ESI-MS spectra)

**Fig. S6:**
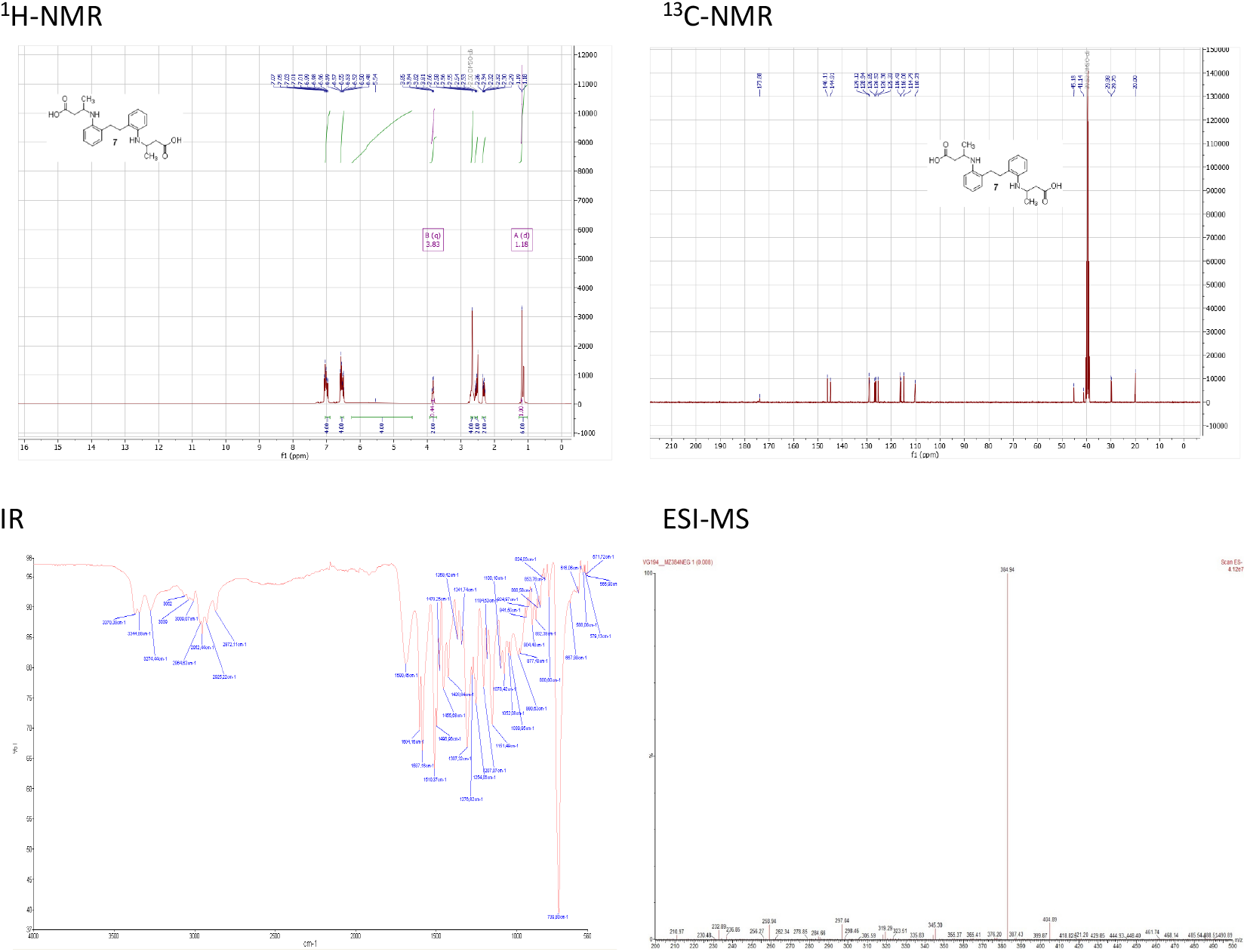
Characterization of compound 7 (BG-194) after chemical synthesis. (^1^H-NMR, ^13^C-NMR IR and ESI-MS spectra)

